# Development and Characterization of a Weaned Pig Model of Shiga Toxin–Producing *E.* coli-Induced Gastrointestinal Disease

**DOI:** 10.1101/2021.08.26.457881

**Authors:** Justin X. Boeckman, Sarah Sprayberry, Abby Korn, Jan S. Suchodolski, Chad Paulk, Kenneth Genovese, Raquel R. Rech, Paula R. Giaretta, Anna Blick, Todd Callaway, Jason J. Gill

**Affiliations:** Department of Animal Science, Texas A&M University, College Station, TX USA; Department of Small Animal Clinical Sciences, Gastrointestinal Laboratory, College of Veterinary Medicine & Biomedical Sciences, Texas A&M University, College Station, TX, USA; Department of Grain Science and Industry, College of Agriculture, Kansas State University, Manhattan, KS, USA; USDA-ARS, Food and Feed Safety Research Unit, College Station, TX USA; Department of Veterinary Pathobiology, College of Veterinary Medicine and Biomedical Sciences, Texas A&M University, College Station, Texas, USA; School of Veterinary Medicine, Universidade Federal de Minas Gerais, Belo Horizonte, Brazil; Department of Animal and Dairy Science, University of Georgia, Athens, GA USA; Center for Phage Technology, Texas A&M University, College Station, TX USA

## Abstract

Post-weaning enteropathies in swine caused by pathogenic *E. coli*, such as post-weaning diarrhea (PWD) or edema disease (ED), remain a significant problem for the swine industry. Reduction in the use of antibiotics over concerns of antibiotic resistance and public health concerns, necessitate the evaluation of effective antibiotic alternatives to prevent significant loss of livestock and/or reductions in swine growth performance. For this purpose, an appropriate piglet model of enterotoxigenic *E. coli* enteropathy is required. In this study, we attempted to induce clinical signs of post-weaning disease in a piglet model using a one-time acute or lower daily chronic dose of a Shiga toxin–producing and enterotoxigenic *E. coli* strain. The induced disease state was monitored by determining fecal shedding and colonization of the challenge strain, animal growth performance, cytokine levels, fecal calprotectin, histology, fecal metabolomics, and fecal microbiome shifts. The most informative analyses were colonization and shedding of the pathogen, serum cytokines, metabolomics, and targeted metagenomics to determine dysbiosis. Histopathological changes of the gastrointestinal (GI) tract and tight junction leakage as measured by fecal calprotectin concentrations were not observed. Chronic dosing was similar to the acute regimen suggesting that a high dose of pathogen, as used in many studies, may not be necessary. The piglet disease model presented here can be used to evaluate alternative PWD treatment options. Furthermore, this relatively mild disease model presented here may be informative for modeling human chronic gastrointestinal diseases, such as inflammatory bowel disease, which otherwise require invasive procedures for study.

**Importance:** Post-weaning diarrhea remains a significant problem in swine production and appropriate models of pathogenesis are needed to test alternative treatment options. In this study, we present an *E. coli* induced piglet model for post-weaning diarrhea, and also explore its translational potential as a model for human intestinal inflammation. Our study here presents two firsts to our knowledge. 1) The first simultaneous analysis of the intestinal microbiome and metabolome through fecal sampling of piglets challenged with Shiga toxin–producing *E. coli*. This is valuable given the limited metabolomics data from swine in various disease states. 2) A comparison of the clinical signs caused by a daily chronic vs one-time dosing regimen of *E. coli*. This comparison is key as infection by pathogenic *E. coli* in real-world settings likely occurs from chronic exposure to contaminated food, water, or environment rather than the highly concentrated dose of pathogen that is commonly given in the literature.

## Introduction

Post-weaning diarrhea (PWD) and edema disease (ED) following the weaning period in piglets remain significant problems for the swine industry and can result in significant economic losses (1-3). PWD is characterized by diarrhea which can lead to severe dehydration, emaciation, and death. While ED of swine is characterized by submucosa edemas of the stomach and mesocolon resulting in swelling of eyelids, forehead, and in some cases hemorrhagic gastroenteritis leading to eventual death (2). Pathogenic *Escherichia coli* is the primary cause of these diseases in swine, and the transitionary period of weaning leaves piglets susceptible to infection by pathogenic strains of *E. coli* (3, 4). While PWD and ED are generally caused by enterotoxigenic *E. coli* (ETEC) and Shiga toxin–producing *E. coli* (STEC), respectively, they affect similarly aged pigs and there can be considerable crossover between serotypes and associated virulence factors. PWD ETEC are primarily associated with *E. coli* producing heat-stable and/or heat-labile enterotoxin, while ED STEC are associated with Shiga toxin, primarily the 2e subtype (Stx2e), producing strains, which can be expressed with or without other enterotoxins (5). The antibiotic colistin has been the classical treatment for pathogenic *E. coli* in swine, however given concerns over antibiotic resistance, alternative treatment options should be explored (1). To evaluate alternative treatment options, a comprehensive ETEC/STEC model of pathogenesis in swine is necessary to evaluate efficacy of alternative treatments to antibiotics. Furthermore, such an ETEC/STEC induced swine model of gastroenteritis could also serve as model for ETEC/STEC induced enteropathy in humans.

Swine are used widely in biomedical research as a proxy for humans, due to similar physiology, immune systems, intestinal permeability, and intestinal enzymatic profiles. Weaned pigs represent a possible model for human ETEC or STEC infections as porcine gastrointestinal anatomy, immune response, and ETEC clinical signs closely mimic that of humans (6). Swine inoculated with ETEC experience sloughing of intestinal villi, increased crypt depths, and scours (7). It has been shown that ETEC infections in weanling pigs can be caused by a single dose of approximately 10^9^ CFU (8, 9). However, this high acute single dose most likely does not accurately represent the real-world scenario of PWD or ED in which piglets are more likely initially infected by chronic exposure to lower doses of *E. coli* as ETEC/STEC can be found in contaminated feed, water, soil, and elsewhere in the barn environment (2). The objective of the present study was to develop and characterize a Shiga toxin–producing *E. coli* induced weaned swine model of PWD/ED. Given the *E. coli* strain used in this work encodes heat-labile enterotoxin IIA (LT-IIA), heat-stable enterotoxin II (STIIB), as well Shiga toxin (Stx2e) it will henceforth be referred to simply as an STEC strain, despite it technically classifying as both an ETEC and STEC. We also sought to evaluate differences in dosing regimens, comparing a one-time high acute dose to a lower daily chronic dose of STEC. To our knowledge, this is the first reported comparison on the effects of a one-time high acute dose vs a lower chronic daily dose in an animal model. Furthermore, comparing the single- or repeated-dose models in swine is critical to being able to evaluate the piglet model as potential model for human ETEC or STEC induced enteropathies, particularly for the study of chronic inflammatory gastrointestinal disorders.

## Results and Discussion

### Growth performance

Pigs used as an experimental model for enteric enteropathy were challenged with a spontaneous nalidixic acid-resistant mutant of *Escherichia coli* strain NCDC 62-57 (ATCC 23545) referred to hereafter simply as *E. coli* 62-57nal in either a single acute high-titer dose (∼10^9^ CFU), or in a series of daily lower-dose challenges (∼10^7^-10^8^ CFU). All pigs were held for two days prior to the start of the trial and were asymptomatic for gastroenteritis. Additionally, pigs were not colonized by organisms capable of forming colonies on MacConkey amended with 50 µg/ml nalidixic acid (MacConkey+nal), and no endogenous phage infecting *E. coli* 62-57nal were identified. Thirty-six presumptive coliform colonies from pooled fecal samples plated on MacConkey agar (0 µg/ml Nal; three colonies per pen) were also tested by PCR for the presence of Shiga toxin type 1 (Stx1), Shiga toxin type 2 (Stx2), heat-stable enterotoxin I (ST1), heat-stable enterotoxin II (ST2) and heat-labile toxin (LTI). All colonies were negative for Stx1, Stx2, ST2 and LTI, but three colonies were positive for ST1. Presence of ST1 gene alone is not a strong predictor of ability to cause disease (8, 10, 11) and pigs were asymptotic, so all animals were retained in the study.

In general, pigs administered *E. coli* 62-57nal via both the acute and chronic dosing regimens presented similar clinical signs with the majority of pens developing scours by day 2 and continuing through day 6. Control pens had visibly soft feces on day 5 and 6 with a single incident of scours on day 6, however the animals in control pens remained visibly healthy throughout the trial period. Additionally, the control pen with the incidence of scours was culture negative for the inoculated *E. coli* 62-57nal throughout the trial, so scours may have been induced by stress or other native microbiota. There was no evidence of difference for overall average daily gain (ADG), average daily feed intake (ADFI), and gain:feed (G:F) of the different treatment groups (P > 0.184, Table 1). However, there were numerical differences between pigs fed the treatments, suggesting that the modest number of replicates and the inherently high post weaning variability in performance were responsible for the failure to detect significant differences in growth performance. This lack of evidence for significant growth differences is similar to previously reported results (12). Pigs administered the acute and chronic dose of *E. coli* 62-57nal had a 54.7% and 14.9% reduction in ADG compared to the control pigs, respectively (Table 1). The control group had the lowest ADFI among the three treatments with acute and chronic dosing regimens increasing feed intake by 17.3% and 29.95%. These findings are in agreement with previous work that showed a 24% decrease in control pigs ADFI compared to the pigs inoculated with ETEC O149 on d 3 to d 6 (9). Madec et al. (13) had similar results with a decrease in weight of weaned piglets inoculated with pathogenic *E. coli* expressing K88 fimbriae from day 0 to day 2 which then recovered by day 9 of the trial. In this study, the acute challenge group had the poorest mean G:F conversion with the control group having the highest mean feed efficiency. Piglets experiencing PWD have been reported to exhibit reduced weight gains (3, 14), however statistically significant reductions in weight performance were not observed, perhaps due to the relatively brief duration of the trial or small sample sizes.

**Table 1.** Effects of ETEC 23545 treatment on nursery pig performance.

Figures and Table Captions
| Table 1. Effects of ETEC 23545 treatment on nursery pig performance |  |  |  |  |  |
| --- | --- | --- | --- | --- | --- |
|  | Treatment |  |  |  |  |
| Item | Control | Acute | Chronic | SEM | Probability, P < |
| **d 0 to 6** |  |  |  |  |  |
| **ADG, g** | 263 | 119 | 225 | 55.93 | 0.224 |
| **ADFI, g** | 179 | 210 | 232 | 43.05 | 0.688 |
| **G:F** | 2.02 | 0.66 | 0.89 | 0.722 | 0.184 |

A total of 24 weaned barrows (initial BW approximately 14 lb; Landrace × Large White) were used in a 7 d study. Pigs were housed 2 pigs per pen with a total of 4 pens per treatment. Treatments refer to the following: Control= Pigs administered a sham challenge of PBS each day, Acute = pigs administered approximately $10^9$ CFU of *E. coli* 62-57nal on d 1 then sham
challenged for the remaining time, Chronic = pigs administered approximately $10^7$ CFU of *E. coli* 62-57nal from d 1 to d 6.

### Bacterial colonization and fecal shedding

The ability of *E. coli* 62-57nal to colonize the gastrointestinal tract of inoculated piglets was determined by measuring colony-forming units recovered from intestinal mucosa, intestinal luminal contents, and in feces. Inoculated strain counts adherent to the mucosal lining were found to be variable, with ∼50% of samples, ranging from 0.21 to 1.71 g of intestinal scraping, yielding counts above the detection limit (5000 CFU/ml of tissue homogenate). Of the samples yielding enumerable colonies, bacterial counts ranged from ∼10^4^ to ∼10^7^ CFU/g in the duodenum, jejunum, ileum, cecum and colon (Fig 1A). Bacterial counts in the cecal and colonic luminal contents, ranging from 0.14 to 11.07g of digesta, were more reliably above the detection limit and ranged from ∼10^3^ to 10^9^ CFU/g, suggesting bacterial proliferation in the unattached population.

**Fig 1.**
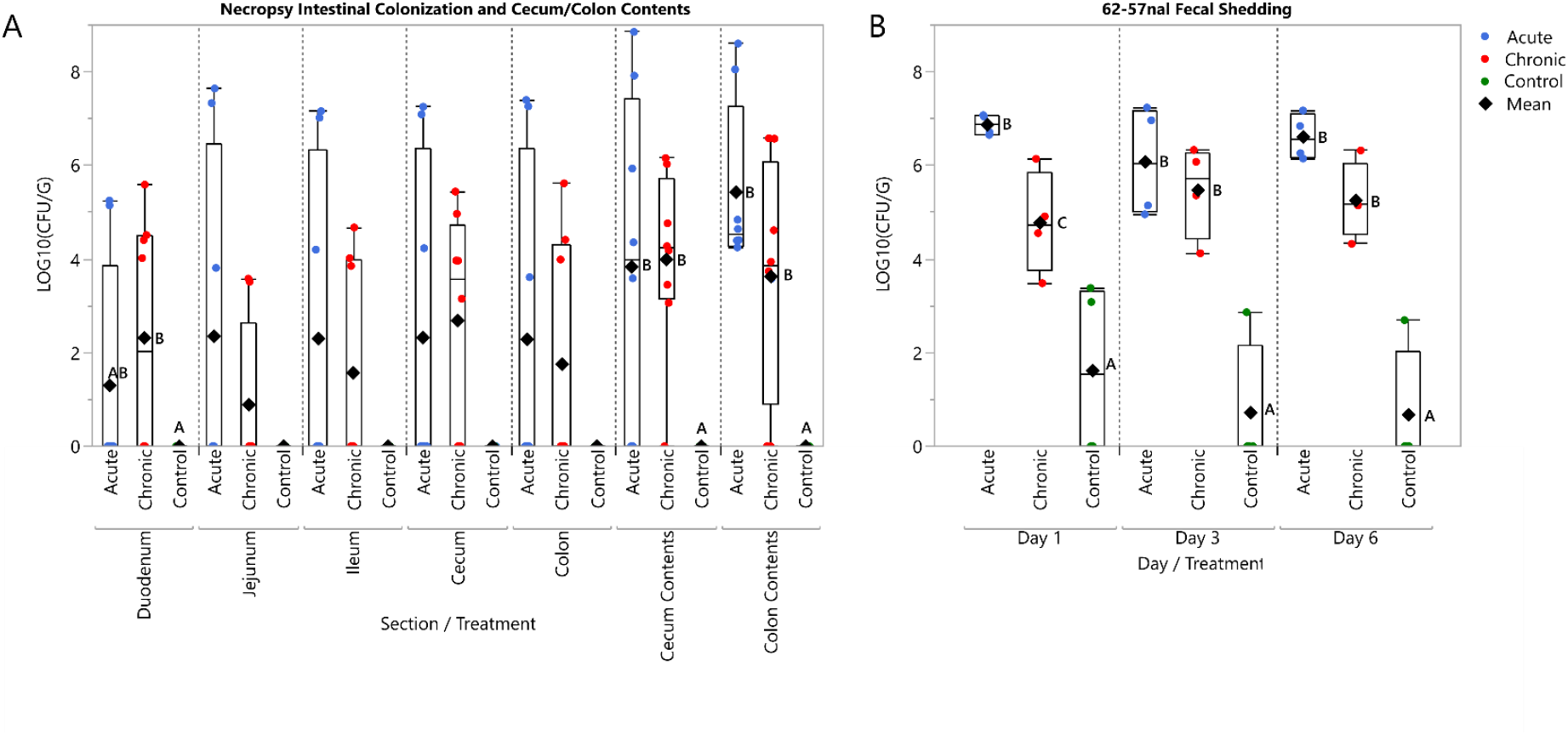
Acute and chronic regimens are sufficient to cause colonization of *E. coli* 62-57nal. **(A.)** At time of necropsy, intestinal scrapings from the duodenum, jejunum, ileum, cecum, and colon were collected and plated for bacterial enumeration. Additionally, cecum and colon contents were collected for direct plating. (**B.)** Fecal samples were collected from each pen on day 1,3, and 6. These samples were serial diluted and plated for direct enumeration of bacteria. LOD∼5×10^2^ CFU/G. **Both (A.) and (B.)** Circles indicate sampling data points, diamonds indicate mean. Sample data was transformed using LOG10. Treatment means grouped by day/section with different letters were significantly different (P<.05)

Acute and chronic treatments had higher prevalence of STEC in feces (∼10^5^ to 10^7^ CFU/g) than control pigs on all sampling days (Fig 1B) Control pens sporadically shed *E. coli* 62-57-nal in the feces at levels near the lower detection limit (500 CFU/g), likely reflecting low levels of pen cross-contamination. Pigs administered the acute STEC dose exhibited significantly higher fecal shedding on day 1 (∼10^7^ CFU/g, P = 0.001) compared to the chronic dose, however there was no statistically significant difference in fecal STEC counts between the acute and chronic treatments after d 1. This result is consistent with other piglet studies which observed peak shedding between 24 and 48 hours post-inoculation (15, 16). Pathogen shedding in the acute group remained high through d 6, indicating that the *E. coli* 62-57nal was able to persistently colonize the gastrointestinal tract of swine.

### Markers of inflammation and intestinal leakage

Infection-induced inflammation is mediated by increased levels of pro-inflammatory cytokines (17). Interleukins 6 and 8 (IL-6 and IL-8) are useful biomarkers since they have been linked to intestinal inflammation (18-21). On d 6 of the study, pigs challenged with STEC had increased (*P* < 0.05) concentrations of serum IL-6 compared to control pigs (Fig 2). However, there was no difference in IL-6 concentrations between acute and chronic treatments. Similar elevations of IL-6 were also observed in the treatment groups of a bacterially induced murine model of chronic intestinal inflammation (22). For concentrations of IL-8, there was a marginally significant overall treatment effect on d 6 (*P* = 0.089): chronic pigs had increased (*P* < 0.05) serum IL-8 concentrations compared to control pigs, and acute dose pigs were intermediate (P =0.5423). Lee *et al*. (23) observed peak serum IL-8 levels in ETEC-challenged piglets between day 0 and 2 which then declined through d 7. This could explain the lower levels of serum IL-8 in the acute challenge group as serum cytokines were measured six days after acute challenge, while the daily chronic challenge may maintain elevated IL-8 concentrations. Increased levels of IL-6 and IL-8 in response to challenge with *E. coli* 62-57nal is in agreement with prior work demonstrating these cytokines as markers of inflammation and infection (22, 23).

**Fig 2.**
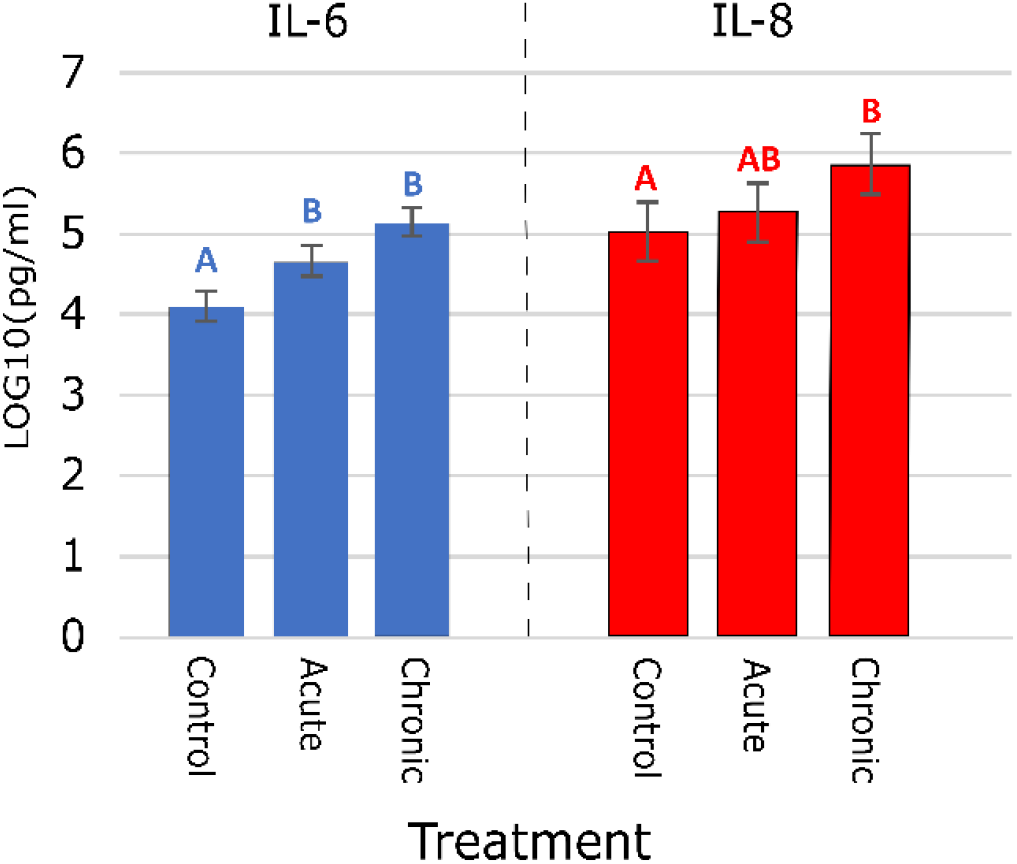
Effect on Serum Cytokines IL-8 and IL-6. On day 6, serum samples were taken from each piglet. Samples were tested by ELISA and data was transformed using Log10. Means with different letters within each cytokine differ significantly (P < 0.05). Error bars represent SEM.

Increased concentrations of fecal calprotectin have been positively correlated with the histological activity of inflammatory bowel disease in humans (24), and serve as a marker of neutrophilic intestinal inflammation (25). Past studies have investigated calprotectin in swine plasma, intestinal lumen, and jejunal mucosa, all of which were found to be correlated with bacterial infection (26). In the present study, there were no significant treatment effects on fecal calprotectin concentration. This is the first study to our knowledge to test fecal calprotectin in pigs inoculated with *E. coli*, and this indicates calprotectin may not be an informative biomarker in this model.

### Villi length and histopathology

Previous studies involving swine challenged with ETEC strains have reported villous atrophy and reductions in crypt depth (27); similar symptoms have also been reported in chronic intestinal enteropathies in humans (28). At time of necropsy, sections were collected to evaluate villi length in the piglet model, but no morphologic changes were observed between treatment groups. General bacterial rod attachment was evaluated by an anatomic pathologist and observed sporadically in all samples with no apparent correlation between rod attachment and direct bacterial plating as only 38% (11/29) of samples with rod attachment tested positive for *E. coli* 62-57nal by direct plating. Villus length in STEC challenged animals did not differ from the controls in the duodenum (P=0.7125), jejunum (P=0.3719), and ileum (P=0.778). Lack of villus blunting may be due to the limited duration of this study. A prior longer-term study (21 d), with a murine model of chronic intestinal inflammation obtained villus blunting through a combination of bacterial challenge and malnutrition (22). Similarly, post-weaning anorexia in piglets has been shown to be associated with reduced villus heights (29). Therefore, given a longer trial period and/or malnourishment, blunting may have been eventually observed in our present model. Histology is also only able to evaluate a tiny fraction of the intestinal tract, so lesions must be broadly distributed throughout the tissue to be detectable by this method. Based on this data, histologic analysis does not appear to be a useful method for evaluating this model.

### Effects on the microbiome by 16S qPCR analysis

To observe any changes of the gut microbiota caused by our acute or chronic dosing treatments, targeted 16S qPCR was performed for select bacterial groups on fecal samples collected from pens at day -1, day 1, day 3, and day 6. Relative abundances obtained were consistent with previous examinations of the piglet microbiome, showing a microbiome dominated by *Bacteroidetes* and *Firmicutes* (30, 31). Overall, the bacterial groups tended to increase relative to control and pre-treatment samples, likely due to natural microbiome succession. A summary of these significant (P < 0.05) or marginally significant (P < 0.10) bacterial group changes at each time point is shown in Table 2. Both acute and chronic STEC doses impacted relative quantities of *E. coli* populations compared to the control. Additionally, both dosing regimens had comparable impacts on microbiome progression. Pretreatment compared to post-treatment samples of the acute dose had the most significant/marginally significant changes with eight of the ten tested bacterial taxa (*Bacteroidetes, Enterococcus, Faecalibacterium, Firmicutes, Lactobacillus, Streptococcus, Fusobacterium*, and Universal) showing increased populations. The chronic treatment showed similar but less dramatic changes, with six of ten taxa (*Bacteroidetes, Enterococcus, Lactobacillus, Streptococcus, E. coli*, and *Ruminococcaceae*) showing increased levels from pre- to post-treatment. The control group showed only two altered bacterial groups, *Enterococcus* and *E. coli*, from pre- to post-treatment. The observed increase for *Enterococcus* and *E. coli* within the control treatment is consistent with the previously reported natural post-weaning piglet microbiome maturation which shows an increase in levels of *Enterococcus* and *Enterobacteriaceae* at 8 days post-weaning (32). The acute dose of *E. coli* had a slightly more pronounced impact on the gut microbiome maturation than the chronic dose, however both acute and chronic treatments were sufficient to cause a detectable dysbiosis.

**Table 2.** qPCR Summary Table. LOGSQ treatment mean compared to Control at each time point (Left) and LOGSQ treatment mean at day 6 compared to its corresponding pretreatment sample at Day-1 (Right). Non-parametric Wilcoxon exact test performed on LOGSQ. Only those significant or marginally significant listed here, significant defined at p<.05 and marginally significant at p<.10. (▲) or (▼) indicate mean LOGSQ are greater than or less than relative to control or Pretreatment. Significant** Marginally Significant*

| Bacterial Groups that change from Control at Day 1 |  | Bacterial Groups that change from Control at Day 3 |  | Bacterial Groups that change from Control at Day 6 |  | Bacterial Groups that change at Day 6 from their corresponding pretreatment sample at Day -1 |  |  |
| --- | --- | --- | --- | --- | --- | --- | --- | --- |
| Acute | Chronic | Acute | Chronic | Acute | Chronic | Acute | Chronic | Control |
| Fusobacterium*<br>▼(.0571) | Fusobacterium*<br>▼(.0571) | E. coli**<br>▲(.0286) | Faecalibacterium**<br>▲(.0286) | Fusobacterium**<br>▲(.0286) |  | Bacteroidetes**<br>▲(.0286) | Bacteroidetes**<br>▲(.0143) | Enterococcus**<br>▲(.0286) |
|  | Enterococcus*<br>▼(.0571) | Streptococcus**<br>▲(.0286) | Universal**<br>▲(.0286) | Ruminococcaceae**<br>▲(.0143) |  | Enterococcus**<br>▲(.0286) | Enterococcus**<br>▲(.0143) | E. coli*<br>▲(.0571) |
|  | E. coli*<br>▲(.0571) |  | Firmicutes*<br>▲(.0571) | Faecalibacterium*<br>▲(.0571) |  | Faecalibacterium**<br>▲(.0286) | Lactobacillus**<br>▲(.0143) |  |
|  |  |  | Bacteroides*<br>▲(.0571) | E. coli*<br>▲(.0571) |  | Lactobacillus**<br>▲(.0286) | Streptococcus**<br>▲(.0286) |  |
|  |  |  | Lactobacillus*<br>▲(.0571) |  |  | Firmicutes**<br>▲(.0286) | E. coli*<br>▲(.0571) |  |
|  |  |  | E. coli*<br>▲(.0571) |  |  | Universal**<br>▲(.0286) | Ruminococcaceae*<br>▼(.0571) |  |
|  |  |  |  |  |  | Streptococcus*<br>▲(.0571) |  |  |
|  |  |  |  |  |  | Fusobacterium*<br>▲(.0571) |  |  |

Principal component analysis (PCA) of 16S qPCR results also provides clear evidence of dysbiosis in STEC-treated groups (Fig 3A). PCA of pre-vs post-treatment samples indicates that by day 6 the acute treatment clearly clustered away from its pretreatment sample, while the chronic day 6 sample showed an intermediate clustering from its pre-treatment sample. In contrast, the control group remained tightly clustered throughout the trial period. This contrast in clustering suggests the microbiome perturbations are induced by the *E. coli* challenge and are not merely normal microbiota progression. The observed relative stability of the control microbiome is consistent with other studies, which reported a microbiome shift immediately after weaning and reached relative stability within 10 days after weaning (31). These findings indicate that both the single acute dose and the chronic lower dose of STEC caused varying degrees of a similar dysbiosis. While in this present study the acute dose of STEC provided a more pronounced microbiome defect, the slight alteration caused by the chronic dose may still be more reflective of chronic subclinical enteropathies.

**Fig 3.**
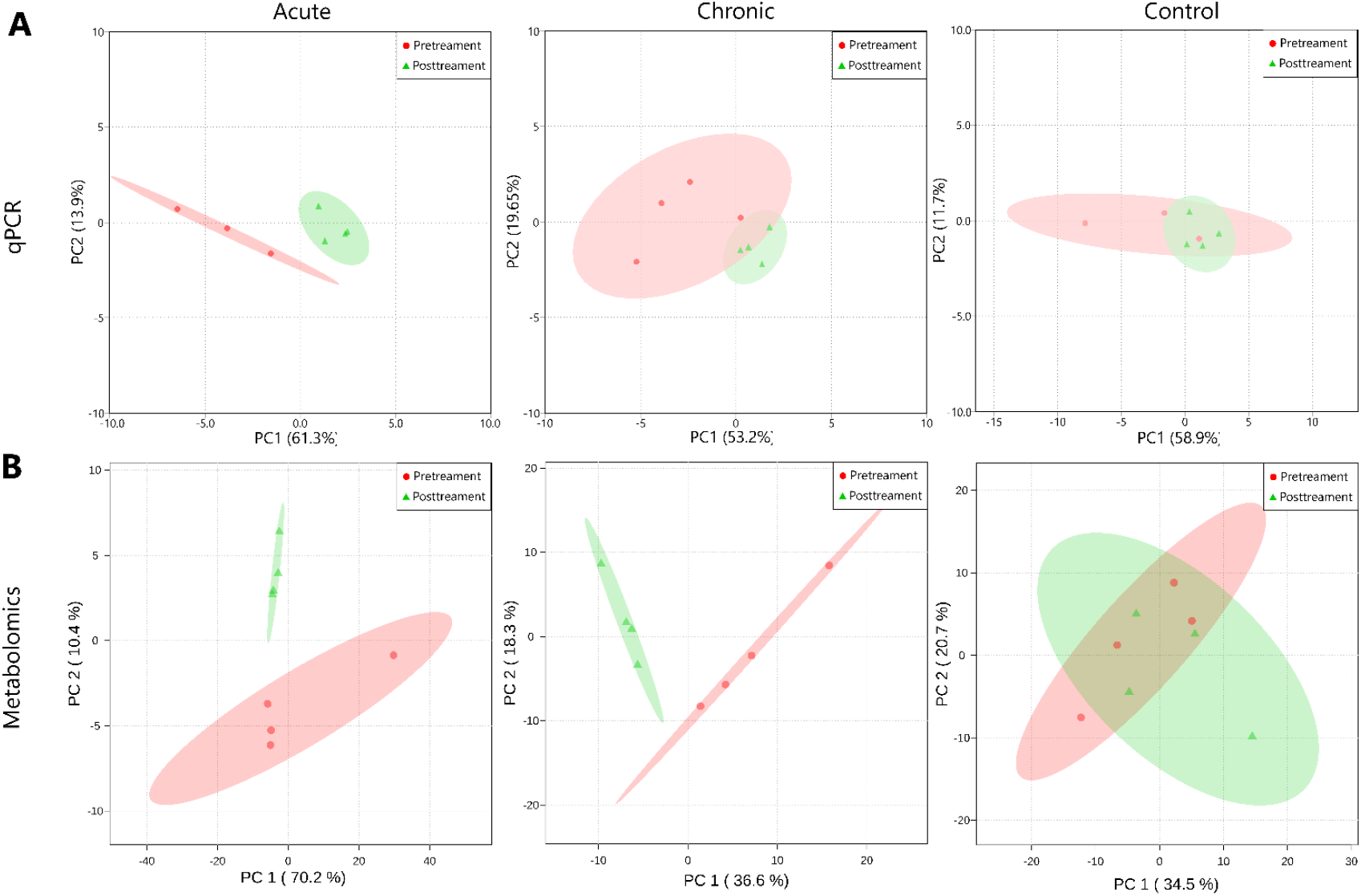
Clustering of treatments by principal component analysis (PCA) of metagenomic and metabolomic results. By the end of the trial period, both chronic and acute treatments are able to be separated from their respective pretreatment samples using qPCR and metabolomic data while the control group remains relatively constant. **(A.)** PCA comparing pre-treatment and post-treatment samples using 16S qPCR of major bacterial taxonomic groups. Red dots indicate pre-treatment samples, green diamonds indicate post-treatment samples, and green/red ellipses represent 95% confidence regions. **(B.)** PCA comparing pre-treatment and post-treatment samples using identifiable fecal metabolites.

### Alterations in the fecal metabolome

To further characterize the differences in disease state caused by acute and chronic STEC challenge, untargeted metabolomics was performed on fecal samples collected pretreatment (d - 1) and days 1, 3 and 6 post-treatment. Metabolite profiles of fecal samples were analyzed by Metaboanalyst (33). Analysis of the identifiable metabolites by PCA clearly distinguished between challenge and control groups (Fig 3B). Similar to the results of microbiome analysis (Fig 3A), the acute and chronic day 6 samples clearly cluster separately from their pre-treatment samples, while the control samples did not separate. The stability of the control group indicates the natural enzymatic, microbial, and structural maturation of the weaned piglet gut (32) was not responsible for the observed shifts in the acute or chronic treatment groups.

As with the 16S qPCR approach, metabolomic comparison of post-treatment samples with their respective pre-treatment samples was more informative when identifying significant changes in individual metabolites. Volcano plots (p-value <0.10 and >2-fold change) were used to identify metabolites that significantly changed following treatment (33) (Fig 4). A full list of metabolites identified by volcano plot is provided in S1 Table. Given the Human Metabolome Database (34) is much more comprehensive, particularly for diseased states, than the Livestock Metabolome Database (35), metabolites identified in this manner were categorized based on the Human Metabolome Database chemical taxonomy. Only four identified metabolites were shared between the chronic and control groups, and ten were common to both the chronic and acute treatment groups (Fig 4). Changes in metabolites in the control group were presumed to be associated with the normal development of the weaned piglet gastrointestinal tract.

**Fig 4.**
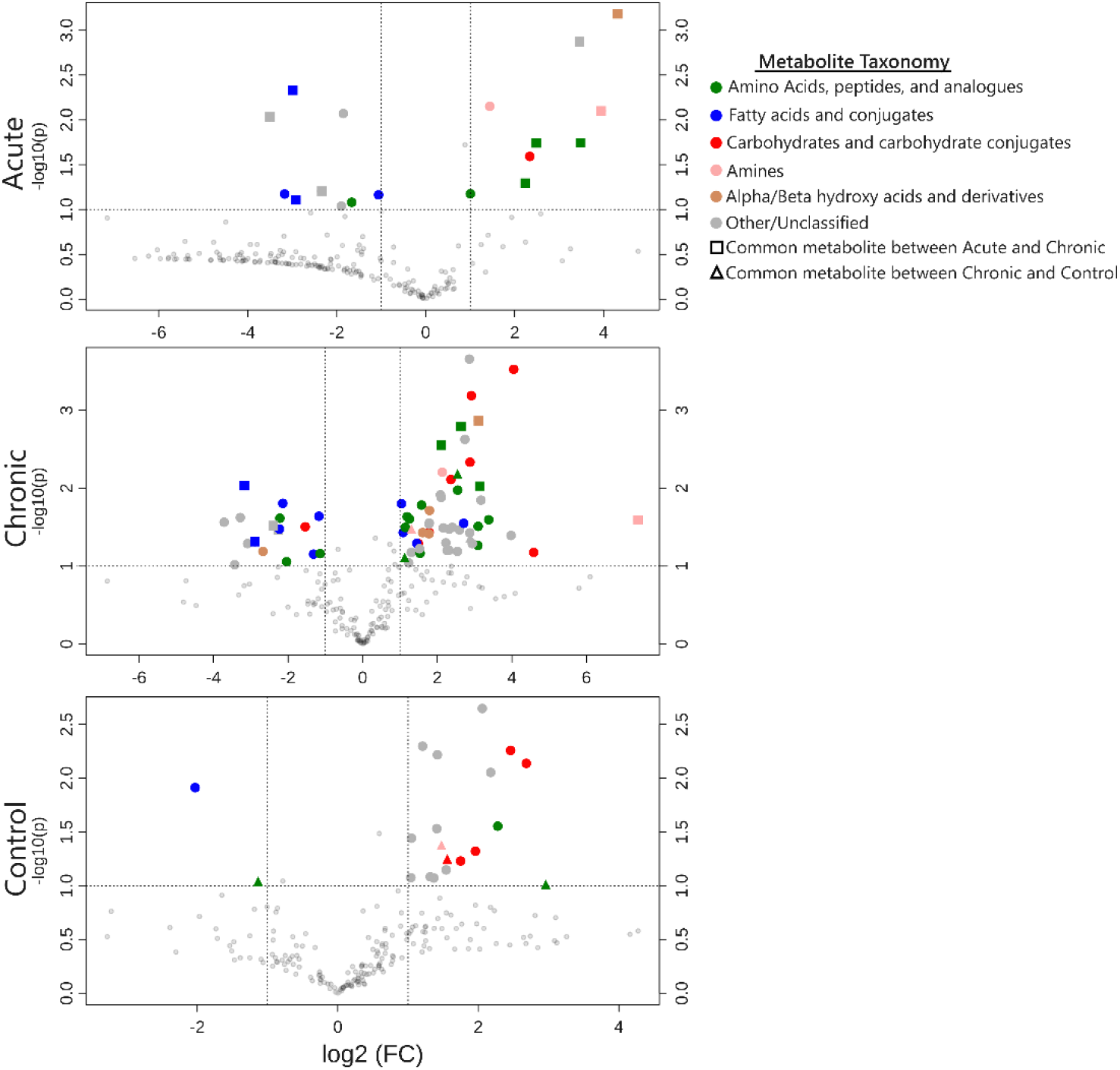
Volcano plots comparing changes in identifiable metabolite profiles in pre- and post-treatment fecal samples. Figures show metabolites that significantly changed from day -1 to day 6 within each STEC dosing regimen. Points represent individual metabolites, which are color-coded based on the Human Metabolomics Database chemical taxonomy (34). Dotted lines indicate significance cutoffs of p-value < 0.1 (Y-axis) and > 2 fold change in abundance (X-axis); points in the upper left and right quadrants of each graph represent metabolites with significant, > 2-fold changes from pre- to post-treatment. Point shape indicates if a metabolite is shared by the acute and chronic treatments (squares) or chronic and control treatments (triangles); circular points indicate metabolites that were either not detected or had non-significant changes in other treatments. Both of the STEC-treated groups exhibited elevated levels of metabolites associated with amino acids and reduced metabolites associated with fatty acids compared to the control group.

Within the chronic and acute treatments, increased levels of amino acid metabolites were identified in the post-treatment samples, including lysine, ornithine, homoserine, histidine, tyramine, beta-alanine, (Fig 4, S1 Table). The increased levels of amino acids and amino acid metabolites in post-treatment fecal samples suggests the STEC treatment led to amino acid malabsorption and/or secretion, likely due to disruption of chemical gradients, inflammation, and microbiome perturbations within the gut caused by STEC treatment. Additionally, metabolites associated with bacterial amino acid degradation, 5-aminovaleric acid and putrescine, were found at increased levels within acute and chronic treatment samples. These have been previously associated with ulcerative colitis (36, 37). The presence of these metabolites is consistent with the model that suggests inflammation caused by STEC treatment induced amino acid malabsorption and subsequent degradation by the resident microbiota. This observation of increased fecal amino acid and amino acid metabolite levels agrees with other studies examining fecal metabolite profiles of humans with inflammatory bowel diseases like ulcerative colitis and Crohn’s disease (38, 39).

Fecal metabolites that were significantly reduced in post-STEC treatment samples were primarily fatty acid metabolites (Fig 4), including stearic acid, myristic acid, and arachidic acid. The levels of lipid-soluble alpha-tocopherol (vitamin E) was also reduced in both treatment groups. The observed depletion of fatty acids within the feces may be indicative of immune system activation, which is supported by our observation of increased levels of serum IL-6 and IL-8. Growing evidence on the importance of “immunometabolism” suggests activated macrophage subtypes and chronically activated T-cells demonstrate increased uptake of fatty acids as they rely more on fatty acid oxidation in order to maintain the high energy levels required to mount an immune response (40, 41). Prior studies examining metabolomic profiles of human inflammatory bowel diseases found pronounced decreases in the levels of short chain fatty acids (SCFA), which are the end products of bacterial fermentation that are absorbed by the large intestine; this presumably signaled a dysbiosis of gut flora (38, 39). In our current study a reduction in the SCFA metabolites butyrate, alpha-ketoglutarate and fumaric acid were observed. Our metabolomic findings indicate both the chronic and acute STEC treatments caused sufficient dysbiosis to statistically distinguish pre- and post-treatment samples (Fig 3B) in large part due to amino acid malabsorption and reduction in fatty acid metabolites (Fig 4), generating metabolomic profiles resembling those of human inflammatory gastrointestinal diseases.

### Whole genome sequencing

To better understand the gene content of *E. coli* 62-57nal that may contribute to its pathogenicity, its genome was sequenced. The genome of *E. coli* 62-57nal was assembled into 378 contigs of greater than 200 bp totaling in 5.6 Mbp length and at an average 45-fold coverage. The entire set of 378 contigs was submitted for sequence typing using SerotypeFinder v1.1 (42) and confirmed to be O138 and H14 as reported previously (43). Analysis of the assembled contigs by BLASTx (>40% identity, E value <10^−5^) against a database of known *E. coli* virulence genes identified a number of genes in *E. coli* 62-57nal associated with pathogenesis, shown in S2 Table. Major identified virulence factors in *E. coli* 62-57nal include hemolysin (*hlyABCD*), Shiga toxin (*stx2e*), an intimin-like adhesin (*fdeC*), heat-stable enterotoxin II (*stiI*), heat-labile enterotoxin IIA (*eltAB*), an iron scavenging cassette (*chuUAVYTWSX*), and the F18ac^+^ fimbrial adhesin. Hemolysin (Hly) is an exotoxin that is associated with many pathogenic strains of *E. coli* (44). Hemolysin is primarily thought to have a role in pathogenesis in extra-intestinal infections, such as those of the urinary tract or septicemia and studies have shown that Hly plays little to no role in clinical signs of diarrhea (45). However recent data, using both in-vivo murine and in-vitro models, show secretion of Hly can disrupt tight-gap junctions and increase colonic mucosal inflammation (46, 47). This inflammation from Hly has been proposed as a contributing factor for ulcerative colitis in humans, a chronic inflammation (46). Consistent with *E. coli* 62-57’s original edema isolation source, an Stx2e Shiga toxin was identified on a putative prophage element. The Stx2e subtype is known to be associated with edema disease of swine (48), however in our present study we did not observe signs of edema in challenged piglets. The primary contributors to the observed scouring phenotype were most likely the identified heat-stable enterotoxin II (STIIB) and heat-labile enterotoxin IIA (LT-IIA). While acting by different modes, both STIIB and LT-IIA have been shown to cause release of water and electrolytes from host membranes, thereby causing diarrhea or scours (11, 49). Heat stable and heat labile enterotoxins are the most common exotoxins that are associated with diarrhea in piglets, present in 72% and 57% of ETEC isolates from piglet scours, respectively (50). *E. coli* 62-57nal also contains a number of virulence factors associated with colonization and survival within the host, including the iron scavenging *chu* genes and various adhesin genes coding for the proteins AIDA-I autotransporter, F18ac^+^ fimbrial adhesin, and the intimin-like FdeC (44, 50, 51). Taken together, the presence of these virulence factors explains the observed diarrhea/scouring and colonization phenotype. Furthermore, the enterotoxin mode of action that induces this diarrhea also disrupts the Na+ gradient (49), which amino acid absorption is dependent upon in the gut (52), likely explaining the increased amino acid levels observed in the fecal metabolomes of ETEC-challenged animals.

## Conclusion

Suitable piglet models exploring the pathogenesis of the *E. coli* pathotypes responsible for common post-weaning diseases like PWD and ED are necessary for evaluating alternative treatment options. Furthermore, appropriate animal models are also imperative for the advancement of knowledge in human disease as well as the development of therapeutic interventions. Rodent models have often been used as disease surrogates, however these models are at a disadvantage when it comes to accurately representing human diseases and syndromes. Swine more accurately resemble humans in anatomy, genetics, and physiology, making them a more appropriate model for human biomedical research (6). The model presented here may serve as a model for evaluation of novel PWD treatments. Additionally, this model could serve as general model for acute or chronic human enteropathies, in which an underlying microbiome dysbiosis is the presumed cause, like ulcerative colitis, Crohn’s disease (53), or environmental enteropathy (28). Many of the differential metabolites identified in this study (Fig. 4) and increases in inflammatory cytokines (Fig. 2) reflect those found in human chronic inflammatory disorders (53, 54). In this study, weaned piglets were challenged with either a single bolus of ∼10^9^ CFU of a pathogenic STIIB+, Stx2e+, and LT-IIA+ *E. coli* strain or daily doses of ∼10^8^ CFU of the same strain. Both chronic and acute treatment groups exhibited significant increases in fecal STEC shedding, intestinal STEC colonization and serum IL-6 levels compared to controls (Fig 1, 2). Furthermore, both treatments induced similar levels of dysbiosis as measured by targeted 16S qPCR and untargeted metabolomics (Fig 3). These findings imply that high acute doses of inoculum, as are often utilized in studies of *E. coli* gastrointestinal disease in pigs, are not necessarily required to establish a disease state, and lower levels of inoculum may represent a comparable disease state important for the study of chronic inflammation. In this study, fecal calprotectin measurements and histological examination of intestinal sections from challenged animals did not indicate any significant differences between control and treatment groups. Given the lack of blunted villi in this work which was previously associated with PWD, future studies employing the model described here may benefit from an extended trial period, as well as controlled changes in animal nutrition, to achieve blunted villi, and if for the evaluation of human disease, possibly the use of a defined humanized microbiota established in germ-free animals.

## Materials and Methods

### Bacterial strains and culture conditions

*Escherichia coli* strain NCDC 62-57 (O138:K81(B):H14) was originally isolated from swine showing clinical signs of porcine edema disease (43). This strain was obtained from the ATCC (ATCC 23545). A spontaneous nalidixic acid-resistant mutant of this strain was isolated and used throughout this work, and will be referred to as strain 62-57nal. This mutation was stable through multiple consecutive transfers. The bacterium was routinely cultured in LB broth (Bacto tryptone 10 g/L, Bacto yeast extract 5 g/L, NaCl 10 g/L) or on LB agar plates (LB broth plus 15 g/L Bacto agar) aerobically at 37°C. Samples obtained from animals challenged with *E. coli* 62-57nal were plated on MacConkey agar (Becton-Dickinson) amended with 50 µg/ml nalidixic acid (MacConkey+nal).

### Whole genome sequencing of E. coli 62-57nal

Total DNA was extracted from an overnight culture of *E. coli* 62-57nal using the Qiagen DNeasy Blood and Tissue Kit following the manufacturer’s specifications for bacterial cells (Qiagen, Cat No. 69504). Isolated genomic DNA was sequenced on the Illumina MiSeq platform using Illumina V2 500 cycle reagent chemistry generating paired-end 250 bp reads. Reads were quality controlled using FastQC (bioinformatics.babraham.ac.uk), FastX Toolkit (hannonlab.cshl.edu), and assembled using SPAdes 3.5.0 (55) at *k-*mer settings of 21,33,55. Contigs <200 bp or with aberrantly low coverage (<8 fold) were filtered from the assembly to yield 378 contigs totaling 5.6 Mbp with ∼45-fold average coverage. The resulting contigs were deposited to Genbank under Bioproject/Accession (PRDF00000000), and underwent automated annotation using the NCBI Prokaryotic Genome Annotation Pipeline (56).

Putative virulence factors were identified based on homology using BLASTx of WGS contigs with a custom database of *E. coli* virulence factors containing *E. coli*-associated proteins contained in mVirDB (57) and characterized *E. coli* virulence factors (44). As a control, the genome of the non-pathogenic lab strain of *E. coli* MG1655 (Accession: GCF_000005845.2) was analyzed against the same database. Hits in common from MG1655 and 62-57nal were excluded based on the presumption that they were part of the non-pathogenic *E. coli* gene repertoire. Protein sequences identified in this initial screening were extracted from the 62-57nal genome and manually investigated using BLASTp and InterProScan to confirm conserved domains were intact and putative gene products were approximately full length (58, 59). The supplied O- and H-antigen serotype provided by ATCC were also confirmed bioinformatically using the SerotypeFinder v1.1 tool located at the Center for Genomic Epidemiology website (42).

### Weaned piglet challenge model of *E. coli* 62-57nal

#### Animals and facilities

All procedures involving animals and their care were approved and monitored by the Animal Care and Use Committee of the USDA Southern Plains Agricultural Research Center (SPARC) and the Texas A&M University Institutional Animal Care and Use Committee. A group of 24 weaned barrows (Landrace × Large White, initial mean BW 6.35 kg) were housed at SPARC in College Station, TX. Pigs were randomly assigned to pens (4.634 m^2^) with 2 barrows per pen that had solid concrete flooring and was equipped with a nipple waterer, rubber mat, and feeder. Pigs were provided *ab libitum* access to water and feed; the diet was a standard phase 1 nursery pig pelleted diet (S3 Table) formulated to meet or exceed the National Research Council (2012) recommended requirements of nutrients.

#### STEC challenge trial

The pigs were held 2 d prior to the start of treatment in order to be pre-screened for endogenous enterotoxigenic *E. coli* (ETEC). Pigs were randomly allotted to one of three treatments: Non-challenged control, acute challenge (a single dose of ∼10^9^ CFU), and chronic challenge (a daily dose of ∼10^7^-10^8^ CFU). Each treatment had a total of 4 pens, 2 pigs per pen, for n=8 pigs total. All pigs were housed in the same barn, with pens separated by empty pens to prevent cross-contamination between treatments. *E. coli* 62-57nal was cultured overnight (16-18 h) in Tryptic Soy Broth (TSB; B-D Bacto) at 37 ºC with aeration. The acute treatment received a single dose of 6 ml of overnight *E. coli* culture on d 1 and the chronic treatment received a daily dosage of 6 ml of a 10-fold dilution in phosphate buffered saline (PBS, Corning Cellgro) starting on d 1 until the termination of the trial. Pigs and feeders were weighed on d 0, 1, 3, and 6 for calculation of average daily gain (ADG), average daily feed intake (ADFI), and gain to feed (G:F). The d 1 collection of weight data, feces, and blood was approximately 12 h after the initial *E. coli* dose was administered. All animals were humanely euthanized and necropsied on d 7 for collection of intestinal scrapings and sections which were used for analysis of *E. coli* colonization and determination of villi blunting, respectively.

### Pre-screening of animals for endogenous phage and ETEC

Prior to the trial initiation (d -1), pigs were screened for endogenous pathogenic *E. coli, E. coli* phages, and enteric bacteria capable of growing in the presence of 50 µg/ml nalidixic acid. Briefly, approximately 1 g of feces from each pen was mixed with 9 ml PBS, vortexed until homogenous, serially diluted (10-fold increments), and plated onto both MacConkey agar plates and MacConkey+nal. A chloroformed aliquot of the first sample dilution was plated onto Tryptic Soy Agar (TSA) plates overlaid with 0.5% top agar (5 g Tryptone, 5 g NaCl, 500 ml dH2O, 0.5% w/v Agar) inoculated with 100 μl from an overnight culture of *E. coli* 62-57nal. MacConkey+nal plates were screened for breakthrough colonies and TSA plates were screened for plaque formation or zones of clearing to determine phage presence. After overnight incubation, 3 colonies were selected from MacConkey plates from each pen and mixed with 150 µl Tris EDTA (TE) buffer. Each sample was boiled for 10 minutes then centrifuged at 8,000 x g for 2 minutes. These colonies were screened via multiplex PCR for: Universal stress protein A (*uspA)*, heat-labile toxin (LTI), heat-stable enterotoxin I (STI), heat-stable enterotoxin II (STII), Shiga toxin type 1 (Stx1) and Shiga toxin type 2 (Stx2) using previously established and validated primers (60, 61). Positive bands of appropriate size were confirmed using individual primer sets and resultant PCR products were visualized on a 1.5% agarose gel with gel red (Biotium).

### Fecal collection and determination of *E. coli* 62-57nal and fecal calprotectin within feces

A representative fecal sample was collected from each pen on d 1, 3, and 6 to determine fecal *E. coli* populations. The samples were collected in individual, sterile 50 ml conical tubes and transported on wet ice to the laboratory. These fecal samples were processed and diluted in the same manner as described above, and were also spot plated (20 µl) on MacConkey agar containing 50 µg/ml nalidixic acid which were incubated for 10-12 h at 37°C to avoid colonies merging within the spots. Fecal calprotectin was determined by a commercially available porcine ELISA kit (MyBioSource, San Diego, CA) with a minimum detection limit of 6.25 ng/ml with an intra-assay CV of less than 15%.

### Blood sampling and serum analysis

Blood samples were collected from 2 pigs per pen on d 6. Blood was collected from the cranial vena cava via a 20 gauge needle and a 10 mL serum vacutainer tube (BD, with clot activator and gel for serum separation). Tubes were inverted and allowed a minimum of 30 minutes to clot. Samples were centrifuged at 1,600 × g for 10 min at 2 °C, and the separated serum samples were stored at -80°C until analysis was performed. Serum concentrations of interleukin 6 (IL-6) and interleukin 8 (IL-8) were determined via porcine ELISA kits (R&D Systems, Minneapolis, MN). The minimum detection for IL-6 was 18.8 pg/ml and 62.5 pg/ml for IL-8. Assays were conducted as outlined by the manufacturer.

### Intestinal sampling and histology

Pigs were humanely euthanized, necropsied, and had samples collected for intestinal histology and *E. coli* populations from the duodenum, jejunum, ileum, cecum, and colon. Segments of the small intestine (duodenum, jejunum, and ileum) and large (cecum and colon) intestine were tied off with plastic zip ties to prevent cross contamination. Adherent bacterial samples were collected by rinsing the intestinal mucosal surface with sterile PBS, scraping a 2-3 cm section of the surface with a glass microscope slide, and then placing the sample into sterile 15 ml conical tubes containing 4.25 g of 2 mm glass beads (Walter Stern Inc.) and 8 ml of sterile PBS. Tissue scraping samples were vortexed for 5 min at 3000 rpm on a platform vortex mixer to homogenize samples. Cecum and distal colon contents were also collected and homogenized by thoroughly vortexing 0.5 - 5 g sample in 25 ml of PBS. Sample homogenates were serially diluted (10-fold increments) in PBS and spot-plated (20 ml) to MacConkey agar with 50 µg/ml nalidixic acid. All bacterial population estimates were normalized to the initial sample wet weight.

For histology, the distal end of each intestinal segment (duodenum, jejunum, ileum, cecum and colon) directly adjacent to the section used for bacterial sampling was collected. Samples were positioned onto a 5.08 cm x 5.08 cm cardboard sections and secured with small clips to prevent tissue curling. Consecutive tissue samples were fixed in Carnoy’s solution (60% ethanol, 30% chloroform, 10% glacial acetic acid) at a 20:1 ratio for 30-45 min and 10% neutral buffered formalin (VWR scientific) at a 10:1 ratio for 24 h, followed by storage in 70% Ethanol until further processing. Tissues were trimmed into longitudinal sections of 5 mm width and embedded into paraffin using standard procedures (62). After processing, 4 µm sections were placed on slides and stained with hematoxylin and eosin and evaluated histologically in a treatment blinded fashion. The slides were analyzed by a board-certified pathologist for rod attachment, presence of inflammation, and morphological changes (i.e., villous blunting, epithelial erosion). A total of 12 pigs, 4 from each treatment, were randomly selected for the evaluation of villus length, and well-oriented and intact villi were measured from the lamina muscularis mucosae layer to villus tip.

### Analysis of fecal metabolites

Fecal samples collected from the floors of pens on d -1, 1, 3 and 6 were lyophilized and sent to the West Coast Metabolomics Center at the University of California Davis for untargeted metabolomic analysis. Untargeted GC-TOF profiling was performed following previously published parameters (63). The resultant dataset is available on Metabolomics Workbench (64), under study number ST001041.

### 16S qPCR

For the purposes of quantifying select bacterial populations, microbial DNA was extracted from 50 mg of lyophilized feces using the Zymo Quick-DNA Fecal/Soil Microbe Kits following the manufacturer’s instructions. Five ng of DNA was used to amplify 16S regions of select bacterial groups using Biorad SsoFast EvaGreen Supermix using reaction conditions described previously (65) and family/genus/species specific primers described previously (65, 66). qPCR data is reported as log_10_ of starting 16S copy number per 5 ng of DNA isolated. Specific primer sets used were for Universal, *Faecalibacterium, Streptococcus, E. coli, Fusobacterium, Firmicutes, Bacteroidetes, Lactobacillus, Ruminoccaceae, and Enterococcus*.

### Statistical analysis

Growth performance along with cytokine and intestinal bacterial population data were analyzed using the PROC MIXED procedure in SAS 9.3 (SAS Inst. Inc., Cary, NC). The model fixed effect was treatment with pen set as a random effect for growth performance, cytokine and intestinal bacterial population data. Fecal samples were collected on a pen basis, therefore pen was not included as a random effect. Calprotectin levels and fecal colony counts were analyzed as repeated measures using the PROC GLIMMIX procedure. Treatment, day, and treatment × day served as fixed effects. Day of collection also served as the repeated measure with pen as the subject. Metabolite data was normalized to sum, mean-centered and divided by the standard deviation of each variable, and analyzed for significant or trending metabolites between treatments using MetaboAnalyst version 4.5 (33). Comparison of qPCR LOGSQ values was carried out using JMP Version 13 (SAS Inst. Inc., Cary, NC.). qPCR treatment means were compared pairwise on a per time point basis; many of the datasets did not pass the Shapiro-Wilk test for normality and were of small sample size, therefore treatments were compared using the nonparametric Wilcoxon Exact Test. qPCR data was also considered using multivariate methods on a pre/post treatment basis using principal component analysis (PCA). Results were considered significant at *P* ≤ 0.05 and marginally significant between *P* > 0.05 and *P* ≤ 0.10.

## Acknowledgements

This work was supported, in whole or in part, by the Bill & Melinda Gates Foundation [OPP1139800]. Under the grant conditions of the Foundation, a Creative Commons Attribution 4.0 Generic License has already been assigned to the Author Accepted Manuscript version that might arise from this submission. This work was additionally supported by Texas A&M AgriLife Research. The authors would like to thank Amanda Blake and So Young Park for assistance with qPCR. We would also like to thank Justin Leavitt, Jacob Chamblee, Jacob Lancaster, Lauren Lessor, Ruoyan Luo, Shayna Smith, Caitlin Older, Jeann Leal de Araujo, Kara Dunmire, Logan Joiner, and Lily Hernandez for their assistance with animal necropsy and sample preparation.

## Data Availability Statement

Data supporting the findings of this study are available within this article and/or supplementary materials, additional supplements can be made available from corresponding author upon reasonable request. Data pertaining to resultant metabolomic analysis is available at Metabolomics Workbench, under study number ST001041. Genomic sequencing data available from Genbank under Bioproject (PRDF00000000).

## Supporting Information

**S1 Table. Metabolites identified by volcano plot**

**S2 Table. *E. coli* 62-57nal virulence genes**

**S3Table. Diet composition (as-fed basis)**

